# Comparative transcriptomic analysis of cassava genotypes under extended photoperiodism across flowering stages

**DOI:** 10.64898/2026.08.06.743231

**Authors:** Michael Landi, Ivan Obare, Trushar Shah, Harriet Okech, Angelyne Abuor, Christine Kanee Mutoni, Morag Ferguson, Andreas Gisel, Leena Tripathi, Samwel Muiruri Kariuki

**Affiliations:** International Institute of Tropical Agriculture (IITA), Nairobi, Kenya; Kenya Plant Health Inspectorate Service (KEPHIS), Nairobi, Kenya; Department of Crops, Faculty of Agriculture, Horticulture and Soils, Egerton University, Njoro, Kenya; Department of Plant Sciences, Kenyatta University, Nairobi, Kenya; International Institute of Tropical Agriculture (IITA), Ibadan, Nigeria; Institute of Biomedical Technologies, CNR, Bari, Italy

## Abstract

Cassava (*Manihot esculenta* Crantz) is a major staple crop across tropical and subtropical regions. Despite advances in genomic selection, delayed, non-flowering, and asynchronous flowering remain key bottlenecks in breeding programs. To better understand the molecular basis of flowering-time variation, we performed RNA sequencing across three genotypes with contrasting flowering phenotypes (early, late, and non-flowering) sampled at three developmental stages under contrasting light regimes in field conditions (natural light and three-hour night-break with white light). Comparative transcriptomic analysis revealed distinct gene expression patterns associated with flowering responses. Genotype comparisons with no light supplementation revealed stage-specific enrichment of biological processes. Light supplementation was associated with changes in the expression of key components of photoperiodic and circadian regulation, as well as pathways involved in flowering-time integration and hormone and sugar-related signaling. These findings suggest that coordinated changes across multiple biological pathways regulate flowering behavior in cassava. The candidate genes and expression patterns reported provide a foundation for functional studies and advance our understanding of molecular mechanisms governing flowering-time regulation in cassava.

## Background & Summary

Flowering is an essential developmental transition in plants, characterized by the shift from vegetative to reproductive growth ^1,2^. This transition is linked to reproductive success, population viability, and adaptation to changing environments. Flowering time influences yield and quality, making it a key agronomic trait in crop improvement and breeding programs ^3,4^. Both endogenous and environmental factors control the timing of flowering ^5-7^. Plants must reach a specific stage of vegetative growth before they can respond to external cues such as light, humidity, and temperature. Flower development is divided into distinct phases: the floral transition, activation of meristem identity genes, specification of floral organ identity, and formation of floral structures. The floral transition phase has been essential for identifying genes involved in flowering time regulation ^8,9^. Studies in model species such as *Arabidopsis thaliana* have identified multiple regulatory pathways controlling flowering time, including photoperiod, temperature, vernalization, and hormone-mediated pathways, with key integrator genes such as *CONSTANS (CO) and FLOWERING LOCUS T (FT)* playing central roles in coordinating these signals ^10-12^. Among these pathways, the photoperiod pathway integrates light signals with the circadian clock to regulate the timing of floral transition.

These regulatory mechanisms are especially relevant in crop species where flowering directly impacts breeding efficiency and yield. Cassava (*Manihot esculenta* Crantz) is a major staple crop cultivated across tropical and subtropical regions, with more than half of global production occurring in Africa. It serves as a primary food source for hundreds of millions of people and plays an important role in food security and rural livelihoods ^13,14^. Cassava is used for food from its storage roots and leaves, processed into starch-based products, and as livestock feed ^14^. While cassava is of considerable socio-economic importance, flowering remains a major breeding constraint because it is delayed, irregular, and often asynchronous. Although cassava is commonly propagated vegetatively, the development of improved cultivars depends on sexual reproduction to enable genetic recombination and the selection of desirable traits ^15^. Many cassava genotypes exhibit delayed, sparse, or absent flowering under ambient field conditions, limiting the ability to generate crosses and reducing the efficiency of breeding programs ^16-18^. As a consequence, breeding cycles are prolonged, often requiring several years due to insufficient seed production and delayed reproductive development ^15,19^. In some cases, agronomically valuable genotypes fail to flower entirely in target environments, further constraining their use in breeding programs ^20^. These challenges highlight the importance of understanding the factors that regulate flowering in cassava to improve breeding efficiency.

At the developmental level, these regulatory processes are reflected in changes in shoot architecture and meristem identity. In cassava, floral initiation occurs when the shoot apical meristem transitions from a vegetative to an inflorescence meristem and is characteristically associated with fork-type branching^16^. Flowering responses in cassava vary across environments, with both temperature and photoperiod influencing flowering behavior in a genotype-dependent manner, including strong interactions in which long-day conditions combined with relatively cool temperatures promote earlier and more abundant flowering ^17^. In contrast to *Arabidopsis thaliana*, where warmer temperatures generally accelerate flowering ^21^, cassava exhibits an opposite response, with warmer conditions tending to delay or suppress flowering, highlighting its sensitivity to environmental cues. Transcriptomic analyses further indicate that flowering regulation in cassava involves multiple environmentally responsive pathways, including those related to photoperiod signaling, sugar metabolism, and hormone regulation, with expression patterns that are conserved with those of known flowering regulatory networks in model species ^22^. Therefore, further understanding of the molecular mechanisms underlying variation in flowering behavior among cassava genotypes would help explain the regulation of flowering time. While previous studies have provided important insights into environmental and transcriptomic regulation, analyses that simultaneously consider multiple genotypes, developmental stages, and environmental conditions could provide a more comprehensive view of flowering control. Such approaches are relevant for improving our understanding of late and non-flowering phenotypes.

Here, we examine gene expression regulation underlying flowering variation across cassava genotypes and developmental stages under contrasting light conditions. Using RNA sequencing, we profiled gene expression in three genotypes with distinct flowering phenotypes, DSC272 (early flowering), DSC120 (late flowering), and DSC196 (non-flowering), across three developmental stages representing the transition from vegetative to reproductive growth. By integrating genotype, developmental stage, and light supplementation within a single experimental framework, this study provides a comparative view of gene expression patterns linked to flowering responses. This approach enables the identification of molecular signatures distinguishing contrasting flowering phenotypes, offering insight into the developmental processes underlying flowering-time variation in cassava.

## Methods

### Plant materials and experimental site

The study was conducted at the Kenya Agricultural and Livestock Research Organization (KALRO) Station, Thika, Kenya. Three cassava (*Manihot esculenta* Crantz) genotypes exhibiting distinct flowering phenologies and varying levels of resistance to Cassava Brown Streak Disease (CBSD) were selected. The genotypes included DSC272, DSC120, and DSC196 (Table 1) ^23^.

**Table 1:** List of clones used in this study and their characteristics.

| Clone | Name | Flowering status | CBSD resistance |
| --- | --- | --- | --- |
| DSC272 | PER597 | Early flowering | Resistant |
| DSC120 | COL144 | Medium to late flowering | Immune |
| DSC196 | ECU 41 | Non-flowering | Immune |

Source<sup>24</sup>.

### Experimental design and growth conditions

Plants were established using a complete randomized block design (CRBD) comprising a light-supplemented treatment and a control group maintained under ambient light. The control and the light treatment experiments were situated 1 km apart within the same KALRO station to prevent inter-plot light contamination. In the treatment blocks, supplemental light was provided by 100-watt white LED floodlights positioned approximately 3 meters above the soil surface to ensure uniform light distribution. In the light treatment, the lights were set so that the night was broken by three hours of supplemental lighting from 10 pm to 1 am. The experimental plots were monitored weekly over a ten-month growth period. Hereafter, the light-supplemented treatment is referred to as “light,” and the control as “no light.”

### Tissue Sampling, RNA isolation, and sequencing

To capture the transition from vegetative growth to reproductive development, tissue samples were harvested at three strategically defined time points: T0 (pre-flowering vegetative stage), T1 (initiation of flowering based-off early-flowering genotype, DSC272), and T2 (full establishment of flowering in early and late-flowering genotypes). Samples were collected from three biological replicates, and to minimize the influence of circadian rhythm-regulated gene expression, all sampling was standardized to occur at approximately 11:00 AM. Harvested tissues were immediately transferred to 50 mL polypropylene tubes, frozen in dry ice for transport, and stored at -80°C to preserve transcriptomic integrity. Total RNA was isolated using a modified sequential CTAB-Trizol protocol optimized for starchy, polyphenol-rich cassava tissues. Initial extraction of total nucleic acids was performed using a CTAB buffer prepared with diethylpyrocarbonate (DEPC)-treated water. Following precipitation and resuspension of the nucleic acid pellet in nuclease-free water, a secondary purification was performed using TRIzol reagent (Thermo Fisher Scientific) for RNA extraction according to the manufacturer’s instructions. This hybrid approach was employed to ensure the removal of complex polysaccharides and secondary metabolites, which frequently interfere with downstream sequencing applications. Sequencing was performed on an Illumina HiSeq 2500 platform, generating 150-bp paired-end reads.

### RNA-seq data processing, quality control, and mapping

Raw sequencing reads were assessed for quality using FastQC (version 0.12.1) (https://www.bioinformatics.babraham.ac.uk/projects/fastqc/). Adapter sequences and low-quality bases from the 5’ and 3’ ends of reads were trimmed using Trim Galore (version 0.6.10) (https://github.com/FelixKrueger/TrimGalore/tree/0.6.10). Clean reads were mapped against the *Manihot esculenta* v8.0 reference genome from Phytozome ^25^ using STAR version 2.7.9a (parameters:--genomeDir, --readFilesIn, --readFilesCommand, -- outFileNamePrefix, --outSAMtype BAM SortedByCoordinate,-- outFilterMismatchNoverReadLmax 0.06,) ^26^. A genome index was first generated using the reference genome sequence and corresponding gene annotation file, with -- genomeSAindexNbases set to 13 based on genome size.

### Normalization of gene expression data and principal component analysis

Gene-level read counts were quantified from the resulting alignments using featureCounts (v1.6.0) ^27^. Genes with low expression counts were filtered before downstream analyses, retaining only those with at least 10 reads in at least 6 of 54 samples. To stabilize variance across a wide range of mean expression values for downstream analyses, we applied DESeq2’s variance-stabilizing transformation (VST). An unsupervised principal component analysis (PCA) was then performed on the VST-transformed data to assess sample clustering and visualize global patterns of variation.

### Differential gene expression analysis

Differential expression analysis was subsequently performed using the DESeq2 package ^28^ with a design that incorporated condition, genotype, time, and their interactions. The analysis included three biological replicates per genotype (three genotypes) at each time point (T0, T1, T2) under two conditions (light supplementation and no light supplementation). Genes were considered differentially expressed at an adjusted p-value < 0.05 and a log_2_ fold change > 1.

### Gene expression clustering and heatmap visualization

To identify patterns of gene expression, variance-stabilized expression values were scaled by gene using the Z-score transformation^29^. Genes were then grouped into six clusters using the k-means algorithm^30^ in R (set.seed = 123). Clusters were ordered based on their mean expression profiles. Heatmaps were generated using the *pheatmap* package, with clustering disabled to preserve the defined cluster structure. Rows represent genes grouped into clusters, while columns represent samples annotated by time point, condition, or genotype comparison.

### Gene ontology analysis and functional annotation

Functional enrichment analysis was performed on significantly differentially expressed genes (DEGs). To enable functional annotation, cassava gene identifiers (Manes IDs) were first mapped to their corresponding *Arabidopsis thaliana* orthologs using the best-match annotation available in the Phytozome v8.0 genome release. Gene Ontology (GO) enrichment analysis was conducted using the R package clusterProfiler ^31^. GO enrichment was performed on the set of *Arabidopsis* orthologs using the org.At.tair.db annotation package, applying the Benjamini-Hochberg (BH) method for multiple testing correction. Additionally, functional annotation of flowering-related genes was curated using the Flor-ID database ^32^ and resources from the Max Planck Institute for Plant Breeding Research (https://www.mpipz.mpg.de/14637/). This enabled targeted investigation of flowering time regulators among the differentially expressed genes.

### Technical Validation

### RNA-seq data quality

We sequenced, on average, 23.37 million paired-end reads per sample, each with a length of 150 bp, generating a total of 70.09 Gb across 54 libraries from three cassava genotypes (DSC296, DSC120, and DSC196) sampled under two conditions (light supplementation and no light supplementation) at three time points (T0, T1, and T2). After trimming, an average of 22.87 million clean reads per sample were retained (Table S1). Read quality was high across all libraries, with mean Phred quality scores exceeding 30 both before and after trimming (Fig. S1), indicating high base-calling accuracy.

### RNA-seq mapping statistics

Clean reads were mapped to the *Manihot esculenta* reference genome (v8.0), with 72.18– 90.00% of reads mapping uniquely (average 83.31%). The average mapped read length ranged from 251 to 253 bp, with a low mismatch rate of 0.49–0.64% (Table 2). These values indicate good mapping quality and overall data reliability for downstream gene expression analysis.

**Table 2:** Summary mapping statistics.

| Sample | Total reads (M) | Aligned (%) | Uniq aligned (%) | Avg. mapped len (bps) | Mismatch rate |
| --- | --- | --- | --- | --- | --- |
| T0_1 | 23.688536 | 84.53 | 81.98 | 252.15 | 0.58 |
| T0_2 | 21.873697 | 80.47 | 78.01 | 252.71 | 0.55 |
| T0_3 | 24.293197 | 86.14 | 83.82 | 252.69 | 0.55 |
| T0_4 | 24.645478 | 84.42 | 81.88 | 252.69 | 0.52 |
| T0_5 | 25.437255 | 84.66 | 82.55 | 252.64 | 0.58 |
| T0_6 | 23.098225 | 74.79 | 72.33 | 252.13 | 0.63 |
| T0_7 | 22.201947 | 80.13 | 77.53 | 252.38 | 0.64 |
| T0_8 | 23.89195 | 83.67 | 81.34 | 252.02 | 0.54 |
| T0_9 | 20.111405 | 82.64 | 80.33 | 252.62 | 0.58 |
| T0_10 | 22.443123 | 92.19 | 89.63 | 253.23 | 0.61 |
| T0_11 | 23.548146 | 77.2 | 74.87 | 251.99 | 0.63 |
| T0_12 | 22.626766 | 74.64 | 72.18 | 252.24 | 0.63 |
| T0_13 | 23.191947 | 80.25 | 78.04 | 252.21 | 0.6 |
| T0_14 | 34.841636 | 87.38 | 85.05 | 253.13 | 0.54 |
| T0_15 | 25.251603 | 79.33 | 77.03 | 252.55 | 0.61 |
| T0_16 | 22.506593 | 76.9 | 74.47 | 252.86 | 0.58 |
| T0_17 | 25.199949 | 87.35 | 84.8 | 252.84 | 0.52 |
| T0_18 | 24.952981 | 84.09 | 81.81 | 253.04 | 0.55 |
| T1_1 | 33.594197 | 82.45 | 80.08 | 252.67 | 0.61 |
| T1_2 | 21.736768 | 86.47 | 84.18 | 252.85 | 0.54 |
| T1_3 | 23.195408 | 89.08 | 86.81 | 253.24 | 0.54 |
| T1_4 | 20.235431 | 88.89 | 86.36 | 253.11 | 0.53 |
| T1_5 | 23.283731 | 86.08 | 83.7 | 252.91 | 0.53 |
| T1_6 | 19.912729 | 87.23 | 84.93 | 252.65 | 0.56 |
| T1_7 | 21.478345 | 91.11 | 88.67 | 253.2 | 0.63 |
| T1_8 | 23.868532 | 85.67 | 83.33 | 253.16 | 0.56 |
| T1_9 | 25.323827 | 85.93 | 83.69 | 253.29 | 0.51 |
| T1_10 | 19.434471 | 89.18 | 86.94 | 253.14 | 0.56 |
| T1_11 | 23.58129 | 86.73 | 84.33 | 253.04 | 0.57 |
| T1_12 | 21.629815 | 84.35 | 82 | 252.68 | 0.62 |
| T1_13 | 23.086712 | 83.65 | 81.44 | 253.09 | 0.56 |
| T1_14 | 20.939863 | 86.27 | 83.94 | 253.28 | 0.6 |
| T1_15 | 22.877464 | 82.61 | 80.31 | 252.95 | 0.55 |
| T1_16 | 22.277372 | 88.19 | 85.85 | 253.02 | 0.54 |
| T1_17 | 21.120945 | 88.26 | 85.75 | 253.13 | 0.61 |
| T1_18 | 19.868339 | 83.35 | 81.06 | 253.2 | 0.52 |
| T2_1 | 19.890828 | 89.61 | 87.26 | 252.98 | 0.53 |
| T2_2 | 22.091826 | 90.47 | 88.06 | 253.48 | 0.51 |
| T2_3 | 22.081306 | 86.03 | 83.63 | 253.17 | 0.54 |
| T2_4 | 22.943031 | 90.29 | 87.88 | 253.05 | 0.51 |
| T2_5 | 20.581188 | 85.74 | 83.39 | 252.74 | 0.55 |
| T2_6 | 23.206738 | 86.36 | 83.74 | 253.27 | 0.52 |
| T2_7 | 20.968638 | 87.37 | 84.93 | 252.73 | 0.55 |
| T2_8 | 23.898505 | 92.47 | 90.1 | 253.45 | 0.49 |
| T2_9 | 22.127759 | 88.14 | 85.87 | 252.88 | 0.5 |
| T2_10 | 20.908915 | 87.87 | 85.37 | 253.38 | 0.61 |
| T2_11 | 23.141043 | 88.42 | 85.97 | 253.12 | 0.54 |
| T2_12 | 21.367819 | 86.94 | 84.42 | 253.21 | 0.6 |
| T2_13 | 21.743683 | 85.59 | 83.33 | 252.46 | 0.54 |
| T2_14 | 22.24253 | 87.24 | 84.97 | 252.89 | 0.5 |
| T2_15 | 19.498323 | 89.79 | 87.44 | 252.99 | 0.51 |
| T2_16 | 21.15823 | 90.93 | 88.59 | 253.26 | 0.53 |
| T2_17 | 23.342918 | 87.18 | 84.94 | 252.92 | 0.52 |
| T2_18 | 22.618843 | 90 | 87.58 | 253.18 | 0.5 |

Unsupervised PCA of variance-stabilized expression data revealed separation of samples along PC1 and PC2, which explained 26.7% and 11.7% of the total variance, respectively (Fig. S2). Samples were partially separated by time point, with T2 distinct from T0 and T1 along PC1, while T0 and T1 showed overlap.

### Genotype-specific gene expression under no light conditions

Differential expression analysis was performed to compare transcriptional differences between genotypes under no-light conditions across the three developmental time points. Pairwise comparisons were conducted between DSC120 vs DSC272, DSC196 vs DSC272, and DSC120 vs DSC196. At T0, DSC120 vs DSC272 showed the highest number of DEGs (487), with a balanced number of upregulated and downregulated genes (242 up and 245 down). In comparison, DSC196 vs DSC272 and DSC120 vs DSC196 exhibited 222 (119 up, 103 down) and 296 (121 up, 175 down) DEGs, respectively. At T1 and T2, DSC120 vs DSC196 showed the highest number of DEGs, with 355 (233 up, 122 down) and 1093 DEGs (676 up, 417 down), respectively (Fig. 2 a). All significant DEGs for each genotype comparison at each time point are provided in Supplementary Tables S10–12, including their expression status at each comparison. Heatmaps of clustered DEGs revealed structured expression patterns across genotype comparisons at each time point (Fig. 3b–d), with clusters showing consistent differences in expression between comparisons.

Enrichment analysis supported the biological relevance of genotype differences. At T0, enriched terms were associated with developmental processes, including meristem organization, pattern specification, and responses to abiotic stimuli (Fig. 2e). At T2, enrichment was observed for photosynthesis, reactive oxygen species metabolism, fatty acid biosynthesis, and stress-related pathways (Fig. 2f). The complete list of enriched GO terms for each timepoint of the genotypes comparisons is provided in Supplementary Tables S5–S6. No significant GO enrichment was detected at T1. These results indicate time and genotype-specific transcriptional differences, consistent with variation in flowering phenotypes across genotypes.

### Light-responsive gene expression across genotypes and timepoints

To assess the biological significance of the dataset, differential expression analysis was performed to evaluate transcriptional responses to light supplementation across genotypes and time points. Comparisons were conducted between light and no-light conditions for each genotype at three time points, before flowering (T0), initiation of flowering (T1), and established flowering (T2). In genotype DSC272 (early flowering), 289 DEGs were identified (T0: 148 up, 73 down; T1: 14 up, 17 down; T2: 11 up, 26 down). DSC120 (late flowering) showed 385 DEGs, with a higher number of upregulated genes at T0 (86 up, 27 down) and increased differential expression at later time points (T1: 91 DEGs; T2: 181 DEGs), including a higher proportion of downregulated genes at T2. In contrast, DSC196 (non-flowering) exhibited the strongest transcriptional response, with 832 DEGs. At T0, 630 DEGs were identified (479 up, 151 down), followed by reduced but consistent responses at T1 (89 up, 29 down) and T2 (55 up, 29 down) (Fig. 1a). All significant DEGs for each genotype are provided in Supplementary Tables S7–S9, including their expression status at each time point. Heatmaps of clustered DEGs revealed structured gene expression patterns within each genotype, with clusters displaying distinct expression patterns across time points and consistent differences between light and no-light conditions (Fig. 1b,d,f).

**Fig. 1:**
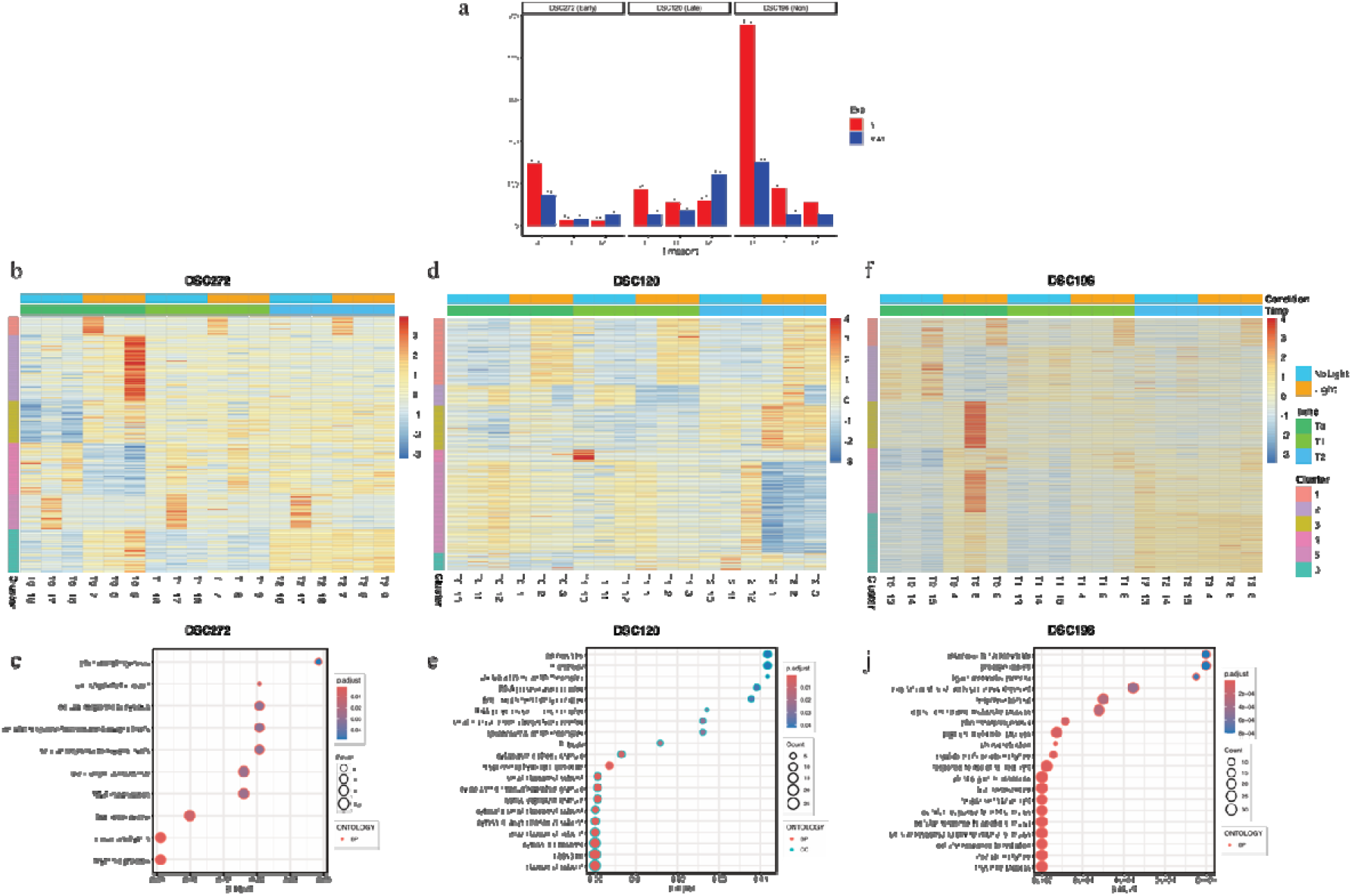
Significant DEGs under light response across genotypes and time points. (a) Number of upregulated (red) and downregulated (blue) DEGs in DSC272, DSC120, and DSC196 under light versus no-light conditions at T0, T1, and T2. (b,d,f) Heatmaps showing clustering of significant DEGs for DSC272, DSC120, and DSC196, respectively. Columns represent samples grouped by condition (Light, NoLight) and time point (T0, T1, T2), while rows represent significant DEGs genes clustered based on expression patterns. (c,e,j) Enriched GO terms of DEGs for DSC272, DSC120, and DSC196, respectively. Dot plots show enriched biological processes, with dot size representing gene count and color indicating adjusted p-values.

**Fig. 2:**
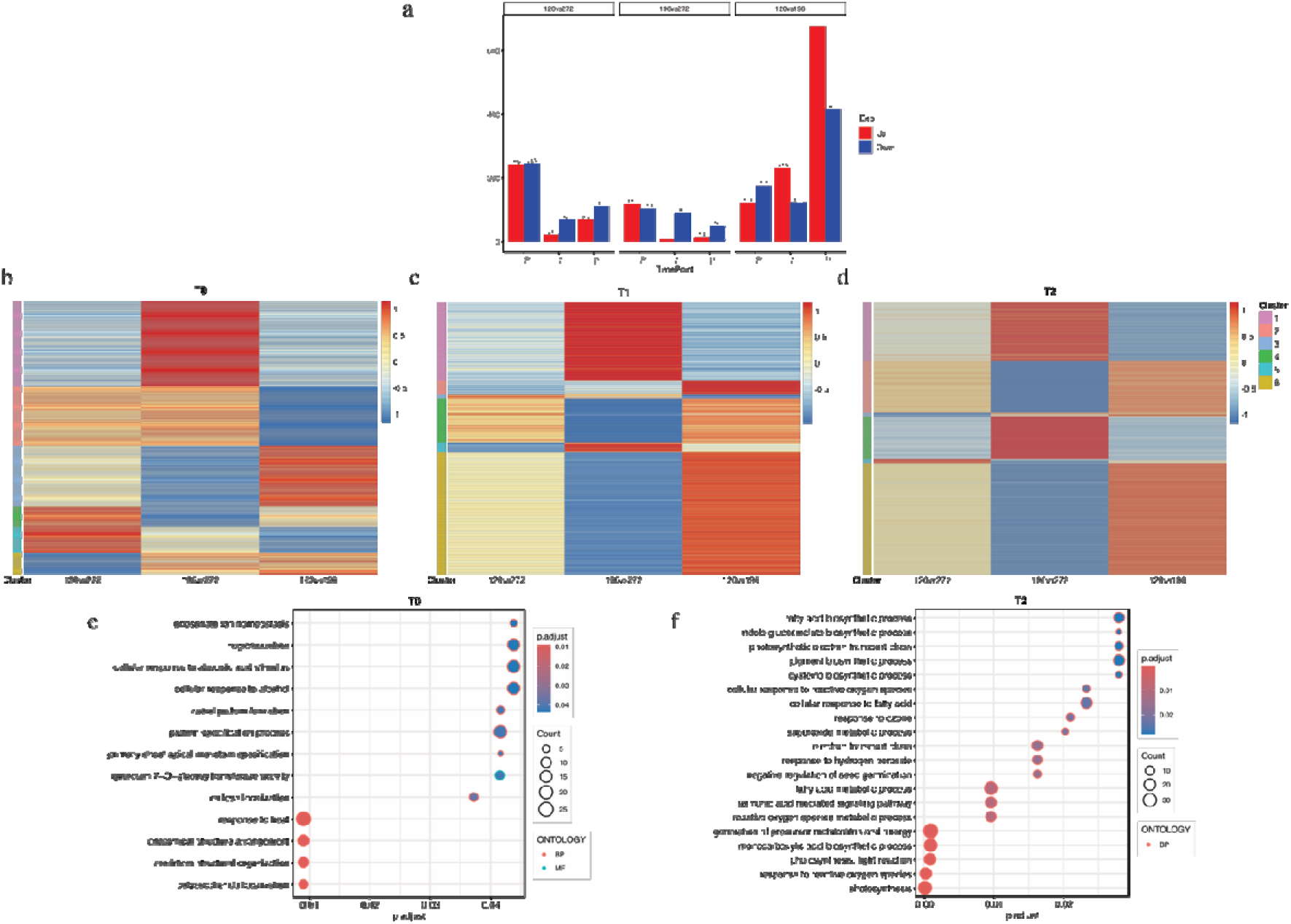
Genotype-specific gene expression under no light condition. (a) Number of upregulated (red) and downregulated (blue) differentially expressed genes (DEGs) for pairwise genotype comparisons (DSC120 vs DSC272, DSC196 vs DSC272, and DSC120 vs DSC196) across time points T0, T1, and T2 under no-light conditions. (b–d) Heatmaps showing clustering of significant DEGs for each time point (T0, T1, and T2). Columns represent genotype comparisons, and rows represent genes grouped into clusters based on expression patterns. (e,f) Enrichment analysis of significant DEGs at T0 and T2, respectively.

Enrichment analysis further supported the dataset’s biological relevance. In DSC272, enriched terms included photomorphogenesis, circadian rhythm, and cellular responses to oxygen levels (Fig. 1c). In DSC120, enrichment was associated with RNA processing and ribonucleoprotein- and ribosome-related processes (Fig. 1e). In DSC196, enriched terms included photoperiodism, response to light stimulus, and photosynthesis-related processes (Fig. 1j). The complete list of enriched GO terms for each genotype is provided in Supplementary Tables S2–S4. These results indicate structured transcriptional responses to light supplementation across genotypes and time points, supporting the overall quality and biological relevance of the dataset. Consistent with these findings, significant DEGs were compared with curated flowering-time regulators and circadian pathway genes. This analysis identified key components of photoperiodic and circadian regulation across all genotypes, including *SPA1-RELATED 3, PSEUDO-RESPONSE REGULATOR 5 (PRR5), LATE ELONGATED HYPOCOTYL (LHY), CONSTANS-LIKE* genes, and *TIME FOR COFFEE*. In addition, genes associated with flowering time integration and meristem identity, such as *FRUITFULL/AGAMOUS-LIKE 8 (FUL/AGL8)* and *APETALA2 (AP2)*, as well as hormone- and sugar-related regulators, including *GA INSENSITIVE DWARF 1B* and *HEXOKINASE 1*, were also identified (Supplementary Tables S13–S15). The presence of these well-characterized flowering regulators supports the validity of the dataset in capturing biologically relevant transcriptional responses to light supplementation.

Overall, this dataset provides a valuable resource for investigating transcriptional responses to light supplementation across cassava genotypes with contrasting flowering phenologies. The identification of key flowering-related and circadian regulatory genes highlights their utility for studying flowering-time variation. The inclusion of late and non-flowering genotypes offers an opportunity to explore genes associated with delayed or absent flowering, which may help inform strategies to induce or improve flowering in these genotypes. Together, these data provide a foundation for future functional studies and may support the identification of candidate genes to improve flowering synchronization and breeding efficiency in cassava.

### Data Records

Raw RNA sequencing data supporting this study are available from the National Center for Biotechnology Information (NCBI) Sequence Read Archive (SRA) database under BioProject accession PRJNA1470176. All supplementary figures, materials and gene expression raw counts are deposited in the Zenodo repository under DOI: https://doi.org/10.5281/zenodo.20759579

## Data availability

Raw RNA sequencing data supporting this study are available from the National Center for Biotechnology Information (NCBI) Sequence Read Archive (SRA) under BioProject accession PRJNA1470176. The corresponding SRA run accessions are SRR38819415, SRR38819416, SRR38819417, SRR38819418, SRR38819419, SRR38819420, SRR38819421, SRR38819422, SRR38819423, SRR38819424, SRR38819425, SRR38819426, SRR38819427, SRR38819428, SRR38819429, SRR38819430, SRR38819431, SRR38819432, SRR38819433, SRR38819434, SRR38819435, SRR38819436, SRR38819437, SRR38819438, SRR38819439, SRR38819440, SRR38819441, SRR38819442, SRR38819443, SRR38819444, SRR38819445, SRR38819446, SRR38819447, SRR38819448, SRR38819449, SRR38819450, SRR38819451, SRR38819452, SRR38819453, SRR38819454, SRR38819455, SRR38819456, SRR38819457, SRR38819458, SRR38819459, SRR38819460, SRR38819461, SRR38819462, SRR38819463, SRR38819464, SRR38819465, SRR38819466, SRR38819467, and SRR38819468.

## Code availability

The software and their versions used for RNA-seq analysis are described in Methods.

## Acknowledgements

We thank the Kenya Agricultural and Livestock Research Organization (KALRO), Thika, Kenya, for providing the field site used in this study. We also acknowledge the International Institute of Tropical Agriculture (IITA) Bioinformatics Unit and team for providing access to high-performance computing (HPC) resources used for data analysis. This work was supported by The Royal Society through the FLAIR Fellowship (FLR/R1/201370) and the FLAIR Collaboration Grant (FCG/R1/211038)

## Authors contributions

S.M.K. conceived and led the project. M.L. performed the data analyses and prepared the first draft of the manuscript. T.S. and A.G. supervised and supported the data analyses and manuscript development. H.O., A.A., C.K.M., and I.O. participated in data collection, field, and laboratory experiments. M.L., I.O., T.S., H.O., A.A., C.K.M., M.F., A.G., L.T., and S.M.K. reviewed and edited the manuscript. All authors read and approved the final manuscript.

## Ethics declarations

**T**he authors declare no competing interests.

